# Frequent paramutation-like features of natural epialleles in tomato

**DOI:** 10.1101/177972

**Authors:** Quentin Gouil, David C. Baulcombe

## Abstract

Freakish and rare or the tip of the iceberg? Both phrases have been used to refer to paramutation, an epigenetic drive that contravenes Mendel’s first law of segregation. Although its underlying mechanisms are beginning to unravel, its understanding relies only on a few examples that may involve transgenes or artificially generated epialleles. By using DNA methylation of introgression lines as an indication of past paramutation, we reveal that the paramutation-like properties of the *H06* locus in hybrids of *Solanum lycopersicum* and a range of tomato relatives and cultivars depend on the timing of sRNA production and conform to an RNA-directed mechanism. In addition, by scanning the methylomes of tomato introgression lines for shared regions of differential methylation that are absent in the *S. lycopersicum* parent, we identify thousands of candidate regions for paramutation-like behaviour. The methylation patterns for a subset of these regions segregate with non Mendelian ratios, consistent with secondary paramutation-like interactions to variable extents depending on the locus. Together these results demonstrate that paramutation-like epigenetic interactions are common for natural epialleles in tomato, but vary in timing and penetrance.

## Introduction

Paramutation is an epigenetic process in plants (including pea, maize, tomato (1)) and animals (worm (2), fruit fly (3), mouse (4)) that is associated with gene silencing. It is unlike other epigenetic mechanisms, however, in that it involves transfer of the silent state from an allele with epigenetic modification to its active homologue. This paramutated allele then becomes silenced and it acquires the ability to silence other active alleles in subsequent generations so that inheritance patterns are non Mendelian (1, 5).

The best characterised examples of paramutation are from maize, at the *b1*, *r1*, and *pl1* loci. Genetic screens have implicated NRPD1 (*rmr6* (6)), the major subunit of Pol IV; the RNA-dependent RNA polymerase RDR2 (*mop1* (7)) and NRPD2a (*mop2*/*rmr7* mutants (8, 9)) the shared subunit of Pol IV and Pol V. These proteins are all required for RNA-directed DNA methylation (RdDM) in which DNA methytransferases are guided to the target sequence in the genome by base pairing of small RNAs (sRNAs). RdDM is also associated with paramutation of the *SULFUREA* locus in tomato (10, 11) and trans-chromosomal DNA methylation in *Arabidopsis thaliana* hybrids (12, 13). Based on these findings the dominant model of paramutation implicates RdDM in the establishment and/or maintenance of the epigenetic mark.

Our interest in paramutation follows from an earlier study of sRNA in tomato lines in which homozygous regions were introgressed from *Solanum pennellii* into *Solanum lycopersicum* (cv. M82) (14). The resulting introgression lines (ILs) each carry many loci at which sRNAs are more abundant than in either parent: they are transgressive (15). To explain these findings we invoked epigenetic mechanisms because, in some instances, there was hypermethylation of the genomic DNA corresponding to the sRNA locus.

In this present study we focus initially on one locus, *H06*, at which there is transgressive sRNA and DNA hypermethylation in multiple introgression lines (15). We were prompted to consider the involvement of a paramutation-like process at *H06* because presence of this epiallele in multiple ILs was not consistent with Mendelian inheritance. The ILs are produced by recurrent backcrossing of the F1 hybrid to the *Solanum lycopersicum* cv. M82 parent so that Mendelian features of the hybrid genome would co-segregate with specific regions of introgressed DNA. At *H06*, however, the segregation must have been independent of any introgressed regions. We could rule out that the anomalous behaviour of *H06* was due to a spontaneous epigenetic change because we could reproducibly recapitulate the transgressive sRNA and DNA hypermethylation at this locus in F2 progeny of the *S. lycopersicum × S. pennelli* cross (15).

An alternative explanation of the *H06* epiallele invoked non-Mendelian inheritance due to a hybrid-induced ‘ paramutation’ that, once triggered, would be inherited in the recurrent backcross progeny independently of any one region in the *S. pennelli* genome. The genetic and molecular tests presented here are fully consistent with that hypothesis and they indicate further that the timing of *H06* paramutation correlates with the production of sRNAs from the paramutagenic epiallele. We also identify other methylated DNA epialleles in the ILs with paramutation-like properties: they are absent from the parental lines, present in multiple ILs independently of a specific introgressed region from *S. pennelli* and they transfer their epigenetic mark to a non-methylated allele following backcrossing to M82. Based on the characterisation of these loci we propose that multiple paramutation-like events occur in the progeny of a *S. lycopersicum* cv. M82 × *Solanum pennellii* cross. These events illustrate how epigenetic effects, of which paramutation is an extreme example, may be induced by hybridisation of divergent genomes. Further characterisation of this epigenetic spectrum will reveal the defining features of paramutation loci and the extent to which hybrid-induced changes to the epigenome can influence transgressive segregation in crop plants and natural evolution.

## Results

### Paramutation at the *H06* locus in multiple lines

The *H06* locus (15) is in the euchromatin of chromosome 8, nine kilobases upstream of two genes in divergent orientation (Fig. 1A). It is unmethylated and lacks small RNAs in M82 but, in IL8-3, it is methylated in all three contexts (CG, CHG and CHH, where H is any nucleotide butG) and produces abundant 24-nt sRNAs (Fig. 1A–C). We refer to this as the *H06*^*IL*^ epiallele. The homologous locus in *Solanum lycopersicum* cv. Micro-Tom and *Solanum pimpinellifolium* has an epigenetic profile similar to *H06*^*IL*^ both in terms of DNA methylation and sRNA production (Fig. 1B and C). In *Solanum pennellii*, however, this locus was distinct from the other species in that there was full methylation at CG and CHG but no sRNAs and only partial CHH methylation in the corresponding region (Fig. 1B and C). This finding is different from our previous report in which the *S. pennellii* locus was described as hypomethylated at all three C contexts (15) (in the aerial part of two-week-old seedlings, whereas here we use only the leaf of a two-week-old seedling). The present data are, however, consistent with the previous finding in that the RdDM features of *H06* mCHH and sRNA are low or absent in *S. pennellii* seedlings.

**Fig. 1.**
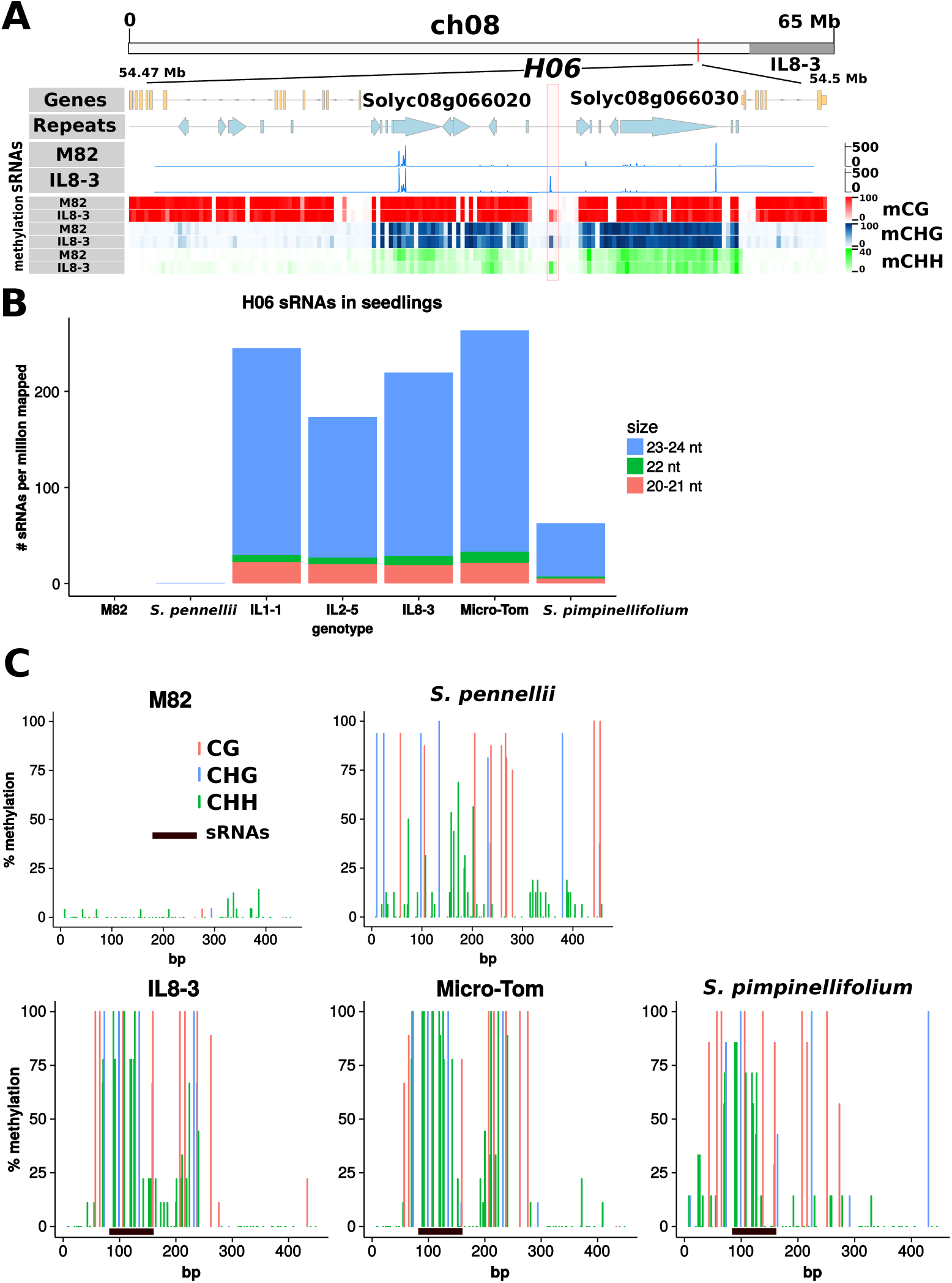
*H06* epialleles. (A) Genomic location of *H06*, sRNAs in M82 and IL8-3 whole seedlings and methylation in seedling leaves. The *H06*^*IL*^ epiallele in IL8-3 was methylated in the CG, CHG and CHH contexts and the source of abundant sRNAs. Neither DNA methylation nor sRNAs were detected in the *H06*^*M*82^ epiallele. (B) High sRNA production at *H06* in seedlings of three introgression lines, the Micro-Tom cultivar and the wild relative *S. pimpinellifolium*. (C) *H06* methylation in leaves of M82, *S. pennellii*, IL8-3, Micro-Tom and *S. pimpinellifolium* determined by Sanger bisulfite sequencing (at least 10 independent clones per genotype). Primer sequences are given in additional file1, and *H06* sequences in additional file 2.

To find out whether the *H06*^*IL*^ has properties consistent with paramutation, we crossed M82 × IL8-3 and monitored the DNA methylation in F1 (M82 × IL8-3), F2 and BC1 (M82 × F1) generations. The DNA was extracted from the leaves of 15-day-old plants and it was assayed with McrBC digestion. The first aim of these tests was to establish first whether the epigenetic mark could be heritably transferred from *H06*^*IL*^ to *H06*^*M*82^ to create an *H06*^*IL′*^ epiallele. The second aim was to find out whether *H06*^*IL′*^ could mediate secondary paramutation and transfer its epigenetic mark to *H06*^*M*82^.

Fourteen of the fifteen F1 plants had highly methylated *H06* DNA in this assay (Fig. 2A) indicating that *H06*^*M*82^ had been converted to *H06*^*IL′*^. The level of methylation in the 30 F2 plants derived from highly methylated F1s was always high, showing that *H06*^*IL*^ and the newly established *H06*^*IL′*^ are normally stable, thus fulfilling the first criterion of paramutation that the epigenetic mark could be transferred from *H06*^*IL*^ to *H06*^*M*82^ to form a stable *H06*^*IL′*^ epiallele. There was, however, instability of *H06*^*IL ′*^ in a single F1 plant and its F2 progeny in which the *H06* DNA was completely unmethylated (Fig. 2A).

**Fig. 2.**
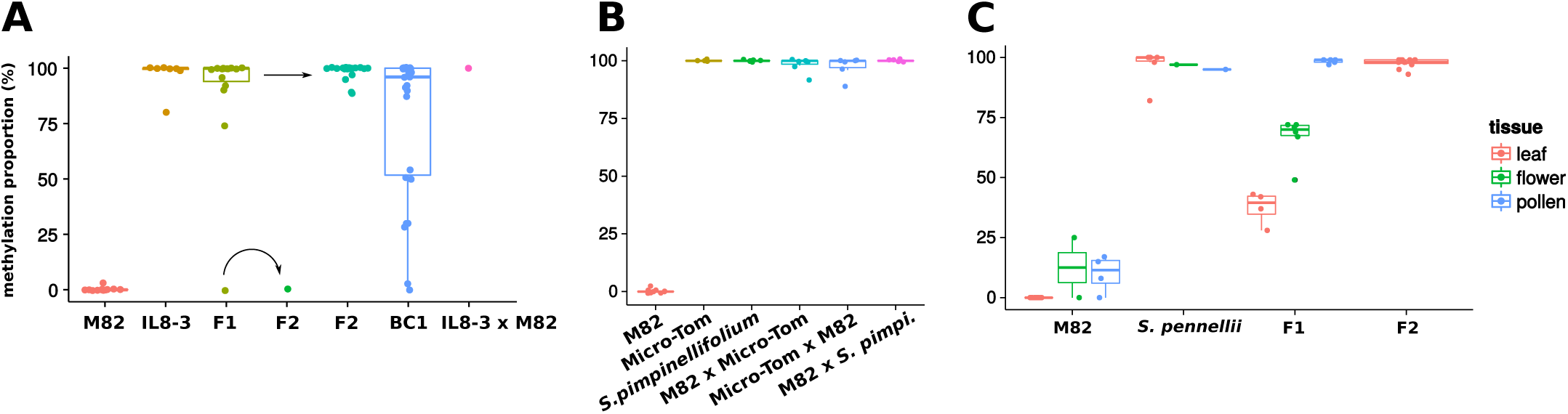
*H06* paramutation assessed by McrBC. McrBC digests methylated DNA, so that comparing amplification of McrBC-treated and non-treated DNA by qPCR reveals the proportion of molecules that were methylated. (A) *H06* McrBC in M82 *×* IL8-3 seedling leaves. Unmethylated *H06*^*M*82^ (n=12), crossed with the methylated *H06*^*IL*^ from IL8-3 (n=7), became methylated in most F1s (14 out of n=15), and remained methylated in the F2 (n=30). Back-crossing the F1 with M82 demonstrated weak paramutagenicity (BC1, n=26, p-value=0.03776, one-sided binomial test). Paramutation also occurred with the IL as the female parent (IL8-3 *×* M82, n=1). (B) *H06* McrBC in M82 *×* Micro-Tom and *S.pimpinellifolium*. F1 seedling leaves are fully methylated (n=6 for each cross), like Micro-Tom and *S.pimpinellifolium* themselves (n=4 and 5 respectively). (C) *H06* McrBC in M82 *× S.pennellii*. The F1 vegetative tissue was half-methylated (n=4) whereas the F1 pollen and the F2 leaves were fully methylated (n=5 and 11 respectively).

The secondary paramutation criterion was tested in the M82 × F1 backcrossed (BC) progeny. If *H06*^*IL*^ but not *H06*^*IL′*^ is paramutagenic then half of the BC1 plants would have fully methylated *H06* DNA and half would have at most 50 percent methylated DNA. In contrast, if both epialleles were paramutagenic, there would be more than half of the plants with full methylation of *H06* DNA. The data (Fig. 2A) are consistent with secondary paramutation because there was an excess of highly methylated plants (18 highly methylated plants, 8 half-methylated or lower, p-value = 0.038, one-sided binomial test). It is likely, however, that some of the *H06*^*IL′*^ or *H06*^*IL*^ alleles in the BC1 were unstable and reverted to *H06*^*M*82^ because five of these BC plants had less than 50 percent methylated DNA at *H06*.

To investigate the paramutagenic properties of the *H06* allele in Micro-Tom, *S. pimpinellifolium* and *Solanum pennellii* we made a further series of crosses with M82 and analysed the F1 progeny. With Micro-Tom and *S. pimpinellifolium* the results were as with IL8-3: all of the *H06* alleles were highly methylated in the F1, irrespective of the direction of the cross (Fig. 2B). In contrast, in M82 × *S. pennellii*, about half of the *H06* alleles were methylated in the 15-day leaves (Fig. 2C). This level increased in flowers and, in pollen and 15d leaves of F2 plants, it was close to 100 percent (Fig. 2C). From these data we conclude that our various *Solanum* genotypes carried three distinct epialleles at *H06*: *H06*^*M*82^, the *S. pennellii* allele referred to as *H06*^*pen*^ and *H06*^*IL*^. The *H06*^*M*82^ allele had neither methylation nor sRNAs. *H06*^*pen*^ had DNA methylation but without the sRNA characteristics of RdDM in leaves of seedlings. This allele triggered an epigenetic change to *H06*^*M*82^ but only after reproductive development in the F1. The third epiallele *H06*^*IL*^ was present in Micro-Tom and *S. pimpinellifolium* in addition to IL8-3, had both abundant sRNAs and DNA methylation in leaves of seedlings, and it could trigger a paramutation-like change to *H06*^*M*82^ without any evident lag. The *H06*^*IL′*^ alleles had the general characteristics of *H06*^*IL*^ but with incomplete penetrance of its effect on *H06*^*M*82^.

### *H06* paramutation timing correlates with sRNA production

To further investigate the interaction of *H06*^*M*82^ with *H06*^*pen*^, we used allele-specific Sanger bisulfite sequencing of *H06* DNA from leaves and pollen of M82, *S. pennellii* and their F1. Consistent with the McrBC results, the allelic methylation of the F1 in the leaves mirrored that of the parents: low methylation of the M82 allele and high methylation of the *S. pennellii* allele (Fig. 3A) at CG and CHG. In pollen, however, the M82 allele of the F1 became hypermethylated in CG, CHG and CHHcontexts (Fig. 3B), whereas pollen in the M82 parent is hypomethylated.

This gain of methylation by the M82 allele in the F1 correlated with an increase in sRNA production at *H06* (Fig. 3C). The *H06* sRNA levels were highest in flowers of *S. pennelli* and the F1, and very low in M82 leaves, flowers and pollen. These levels were, however, much lower than in IL8-3: in the flowers of *S. pennellii* there were 1–2 reads per million mapped whereas in IL8-3 seedlings there were more than 150.

**Fig. 3.**
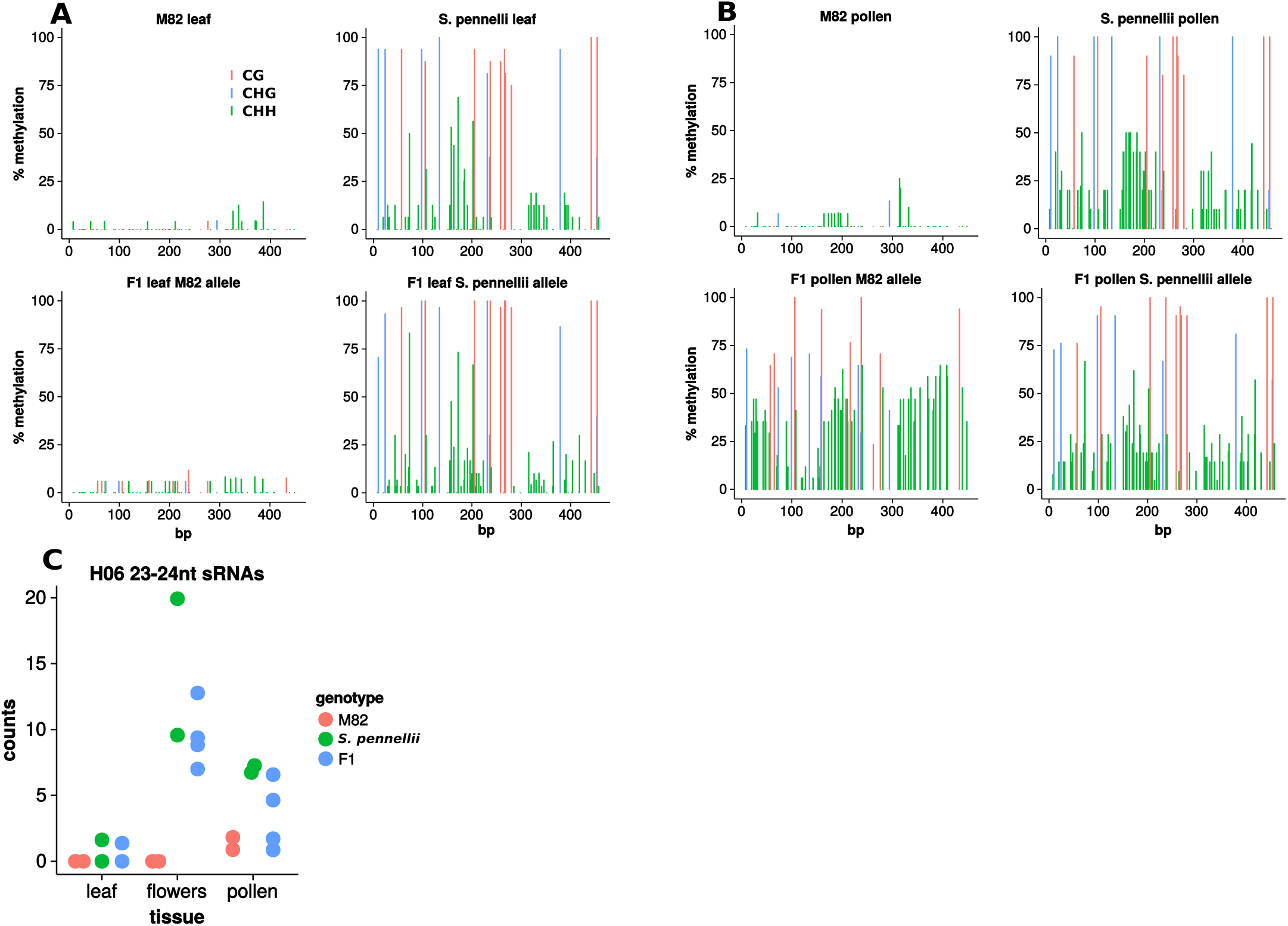
Paramutation in the flowers of the M82 *× S. pennellii* F1. H06 methylation in leaf (A) and pollen (B) of M82, *S. pennellii* and the F1. At least 10 independent Sanger bisulfite clones were used for each plot. (C) *H06* sRNAs are produced in *S. pennellii* and F1 flowers, although at low levels (normalised counts in libraries of about 10 million reads). Two M82, two *S. pennellii*, and four F1 libraries for each tissue.

From these data there is a clear correlation of sRNA with the paramutation-like properties of the various *H06* loci. *H06*^*M*82^ lacks both sRNA and DNA methylation and is paramutable; *H06*^*pen*^ with low levels of sRNAs triggers a delayed paramutation-like process and *H06*^*IL*^, at which the sRNA levels are high, mediates a rapid transfer of the epigenetic marks to *H06*^*M*82^. This correlation of paramutation and sRNA implicates RdDM in the process. Furthermore the changes in both mCHH and sRNA in pollen and flowers suggest that the reproductive phase may be a key developmental stage in the transfer of an epigenetic mark between alleles of *H06*.

### *De novo* methylated *H06* recapitulates paramutation

In principle the association of RdDM with the paramutation-like properties of *H06* could be either a cause or consequence of the transition from *H06*^*M*82^ to *H06*^*IL*^. To test the possibility of a causal role of RNA we used virus-induced gene silencing (VIGS) in which an RNA virus is modified to carry a small host genomic sequence insert. Such RNA viruses may direct DNA methylation of the corresponding genomic DNA of the infected plant (16) and we have previously used this approach to recapitulate the silencing of the *sulf* locus in tomato (11).

To test this system at *H06*, we infected unmethylated M82 plants with a Tobacco Rattle Virus (TRV) containing a 500bp *H06* sequence (TRV-H06, Fig. 4A). Out of 11 successfully infected plants (V_0_ generation) spread across three replicate experiments, only one plant gave rise to methylated progeny (V_1_ generation) (Fig. 4B), with 30% of the plants having high methylation (20/59). None of 6 the control plants infected with unmodified TRV produced methylated progeny (56 tested V_1_ plants) and, with many tens of M82 plants tested over a period of three years, we have never observed spontaneous methylation of *H06* DNA. It is likely therefore that the new epiallele (*H06*^*V IGS*^) was triggered by the TRVH06 VIGS.

**Fig. 4.**
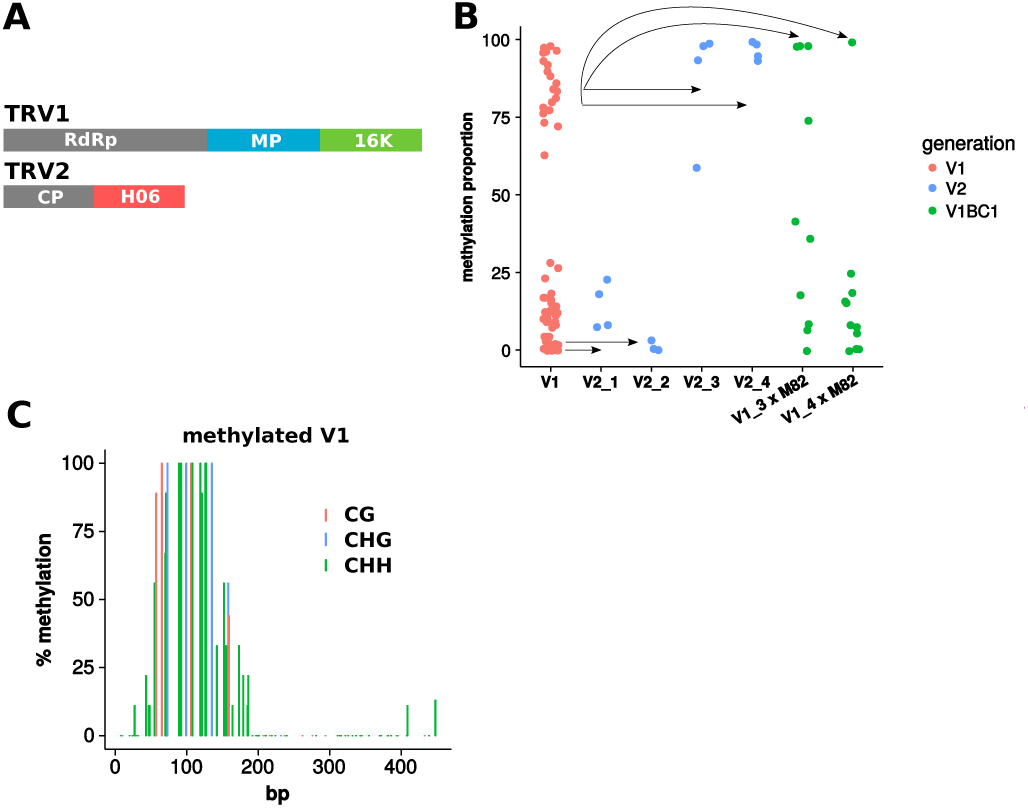
Virus-induced paramutation. (A) Tobacco Rattle Virus modified to contain a 500-bp *H06* fragment. (B) DNA methylation in the progeny of the V_0_ infected plant that gave methylated offspring, assessed by McrBC. V_1_ methylation patterns were stable in the V2 generation, and exhibited weak paramutagenicity in backcrosses to M82 (V_1_ BC_1_) (C) *H06* methylation of a V_1_ plant assessed by Sanger bisulfite sequencing (9 independent clones).

This new epiallele (*H06*^*V IGS*^, Fig. 4C) was distinct from *H06*^*pen*^ and *H06*^*IL*^ (Fig. 1C) in that the hypermethylation was in a restricted region of 200 bp rather than 500 bp. The *H06*^*V IGS*^ epialleles were stable in the V_2_ generation (Fig. 4B). We tested whether this new epiallele was paramutagenic by backcrossing V_1_ plants to M82: if *H06*^*V IGS ′*^ is inherited as a standard Mendelian locus without paramutation, the BC1 plants would have no more than 50 percent of methylated *H06* DNA in an McrBC assay; if however *H06*^*V IGS ′*^ is paramutagenic, some backcrossed plants would exhibit methylation above 50%. Of 21 BC1 plants, 4 had substantially more than 50 percent of methylated *H06* DNA (Fig. 4B), and we conclude that *H06*^*V IGS*^ has weak paramutagenic activity. There were also many plants with less than 50 percent of methylated *H06* in these BC1 plants that is due, presumably, to instability of *H06*^*V IGS ′*^ in the heterozygous condition. The ability to epigenetically modify and confer paramutation properties to *H06* by VIGS indicates that sRNAs could be causal in paramutation.

### Genome-wide ’ paramutation’ in introgression lines?

Is *H06* paramutation an isolated example, or could there be other similar loci in *Solanum* genomes? To address this question we screened the DNA methylome of M82 and three different introgression lines, IL1-1, IL2-5 and IL8-3, in duplicates (summarised in additional file 3). We reasoned that paramutation loci would be differentially methylated compared to M82 in multiple introgression lines. To identify DMRs with this characteristic we compared methylation counts in CG, CHG and CHHof each introgression lines to M82 in 300-bp sliding windows. Each introgression line had 10,000–30,000 DMRs depending on context (CG, CHG and CHH) as compared to M82 (Fig. 5). Only a subset of these DMRs were shared in all three ILs: around 4,600 CG hypermethylated DMRs, 1,800 CG hypomethyated DMRs, 3,600 CHG hypermethylated DMRs, 1,000 CHG hypomethylated DMRs, 262 CHH hypermethylated DMRs and 900 CHH hypomethylated DMRs (Fig. 5 and additional file 4). Of these DMRs there were 25, including *H06*, that were hypermethylated in all three contexts (additional file 5). There were also 10 hypomethylated DMRs in all three contexts (additional file 6).

**Fig. 5.**
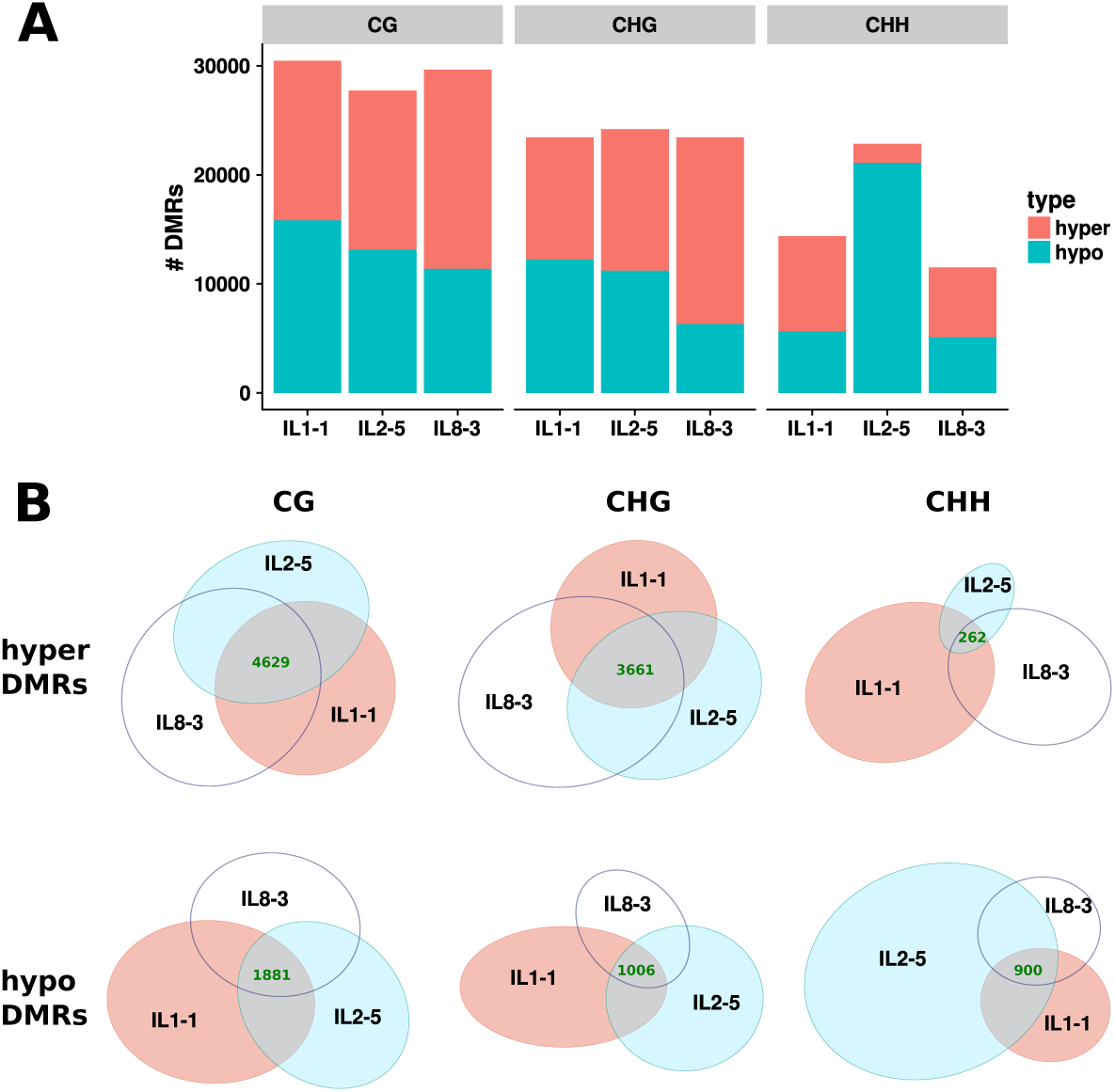
Introgression lines DMRs. (A) Number of DMRs in each introgression line compared to M82 for CG, CHG and CHH contexts. (B) DMR overlap across introgression lines. Hypermethylated DMRs have higher methylation in the introgression lines compared to M82. Area-proportional Venn diagrams.

The RdDM model of paramutation suggests that there would be 24-nt sRNAs at the paramutated loci, as is the case for *H06*. Using seedling sRNA libraries from (15), we identified 134 loci upregulated in the ILs, and 68 down-regulated (Fig. 6A and B). Of the 134 upregulated sRNA loci, 9 overlapped hypermethylated DMRs and 3 overlapped hypomethylated DMRs. In addition to *H06*, one other locus (ch12:62124801–62125100, referred to as Hyper1) had increased sRNAs and hypermethylation in all three contexts in the introgression lines (Fig. S1). This locus and the other hypermethylated DMRs that overlap differential sRNAs are prime candidates for paramutation that would follow the same pattern as *H06*.

**Fig. 6.**
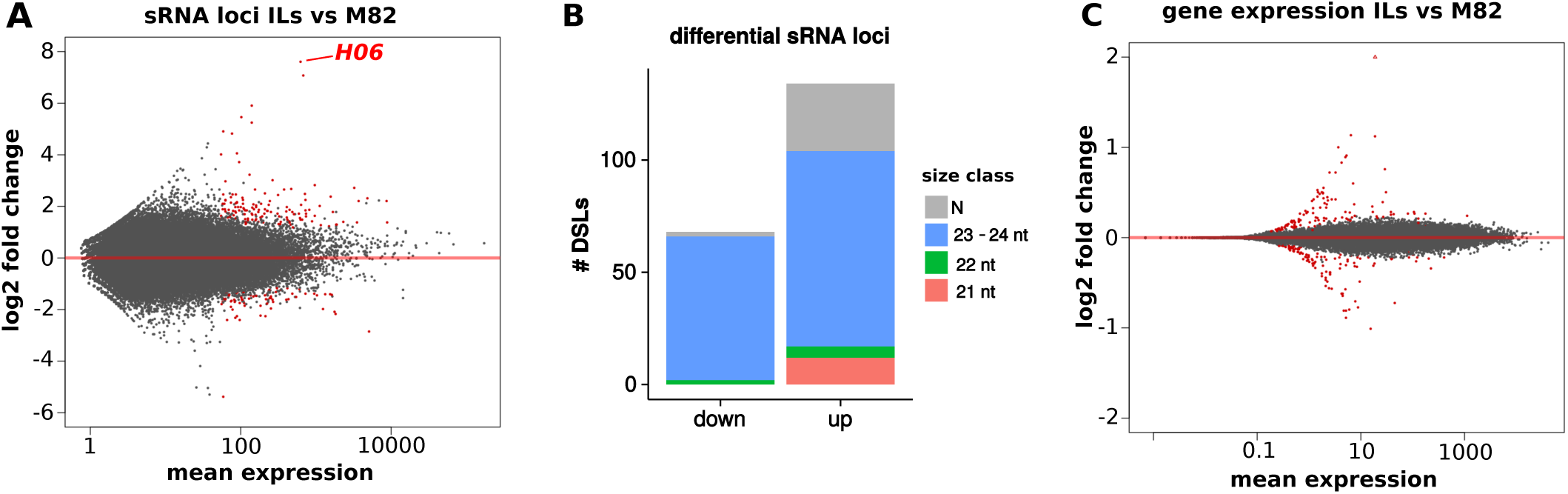
Differential sRNA and gene expression in the introgression lines. (A) Differential analysis of sRNAs between M82 and IL1-1/IL2-5/IL8-3 seedlings. Data from (15). (B) Classification of differential sRNA loci by ShortStack. (C) Differential gene expression between M82 and IL1-1/IL2-5/IL8-3 seedlings. Data from (17). sRNA loci and gene that are differentially expressed (p-adj *<* 0.05) are coloured in red.

DMRs can be associated with differences in gene expression, so we performed differential expression analysis between M82 and the three ILs under scrutiny with data from (17). In this data set for seedlings grown in sun and shade, we found 124 genes that were upregulated in ILs compared to M82, and 108 that were down-regulated (Fig. 6C). The reported log2 fold changes of the differentially expressed genes were modest, suggesting that on/off switches are absent or affect only lowly expressed genes. 10 and 5 of the upregulated genes were overlapped (over their 2 kb promoter or transcribed sequence) by shared hypermethylated and hypomethylated DMRs, respectively, while 4 of the down-regulated genes overlapped hypermethylated DMRs and 7 hypomethylated ones. These genes may be of particular interest to investigate transcriptional and physiological effects of paramutation following a cross between M82 and *S. pennellii*. Although this data set encompasses two environmental conditions (sun and shade), DMRs may affect the transcription of more genes in a tissue- or development-specific manner.

### Paramutation-like activity of DMRs

#### Hypermethylated DMRs

To evaluate the paramutagenic properties of the hypermethylated DMRs, we selected 9 regions based on hypermethylation in multiple cytosine contexts in the three ILs, amenability to McrBC assays, presence of differential sRNAs, and proximity to differentially expressed genes (Hyper1–9, Table 1, Fig. S1). We tested their inheritance in the progeny of M82 × IL8-3 by McrBCqPCR. Mendelian segregation would give 50% methylation in the F1, and in the F2 either 100% (1/4 progeny), 50% (1/2 progeny) or 0% (1/4 progeny) methylation. We tested these DMRs in 10 different F1s and in 33 F2s from three different F1 plants. Any evidence for more than 50% methylation of the F1 alleles or more than 3/4 F2 progeny with 50% or higher methylation would indicate non-Mendelian inheritance and evidence for a paramutation-like mechanism.

**Table 1.** Summary of paramutation candidates and their validation.

| DMR | chr | start | end | type | differential sRNAs | DE nearest gene | evidence for heritable methylation changes |
| --- | --- | --- | --- | --- | --- | --- | --- |
| Hyper1 | ch12 | 62124801 | 62125100 | hyperCG/CHG/CHH | yes | no | strong, incomplete penetrance |
| Hyper2 | ch02 | 48226401 | 48226800 | hyperCG/CHG/CHH | yes | no | strong, incomplete penetrance |
| Hyper3 | ch09 | 61960201 | 61961700 | hyperCG/CHG | no | yes | weaker |
| Hyper4 | ch04 | 26469001 | 26469500 | hyperCG/CHG | no | yes | weaker |
| Hyper5 | ch09 | 61962201 | 61962500 | hyperCG/CHG/CHH | no | yes | no |
| Hyper6 | ch08 | 59748601 | 59748900 | hyperCG | yes | no | no |
| Hyper7 | ch04 | 58749201 | 58749500 | hyperCG/CHG/CHH | yes | no | inconclusive |
| Hyper8 | ch12 | 39437001 | 39437700 | hyperCG/CHG | no | yes | inconclusive |
| Hyper9 | ch04 | 26561001 | 26561500 | hyperCG/CHG | no | yes | no, high variability |
| Hypo1 | ch07 | 12086201 | 12088700 | hypoCG/CHG | no | no | strong, incomplete penetrance |
| Hypo2 | ch06 | 31383601 | 31384100 | hypoCG/CHG | no | no | weaker |
| Hypo3 | ch06 | 47125401 | 47125700 | hypoCG/CHH | yes | yes | weaker |
| Hypo4 | ch03 | 1025801 | 1026100 | hypoCG/CHG | no | no | no |
| Hypo5 | ch09 | 67640601 | 67641100 | hypoCG/CHG | no | no | no |
| Hypo6 | ch05 | 22508201 | 22508700 | hypoCG/CHG | no | no | no, high variability |
| Hypo7 | ch12 | 34481401 | 34482700 | hypoCG/CHG/CHH | no | no | no, high variability |
| Hypo8 | ch03 | 1023801 | 1024100 | hypoCG/CHG | no | no | no, high variability |

According to these criteria, there was evidence of a paramutation-like process at four loci (Hyper1–4, Fig. 7A and Table 1). For three of those (Hyper1–3), some F1s had more than 50% of methylated alleles. At the fourth locus (Hyper4, Fig. 7A) the F1 progeny had approximately 50% of methylated alleles consistent with Mendelian inheritance but more than 3/4 of the F2 progeny had more than 50% methylated alleles (only 3/33 F2s had low methylation, p-value = 0.021, one-sided binomial test). This pattern is compatible with a partial gain of methylation by the M82 allele later in the development of the F1 (as was the case for *H06* in M82 × *S. pennellii*). Of note, Hyper1 corresponds to a region of highly transgressive sRNAs (log2 fold change of 7) that are 24-nt in length (Fig. S1). It is therefore very similar in behaviour to *H06*. As for Hyper3, it is located in the immediate promoter of the Solyc09g064640 gene, which is expressed at lower levels in the introgression lines compared to M82 (Fig. S2).

**Fig. 7.**
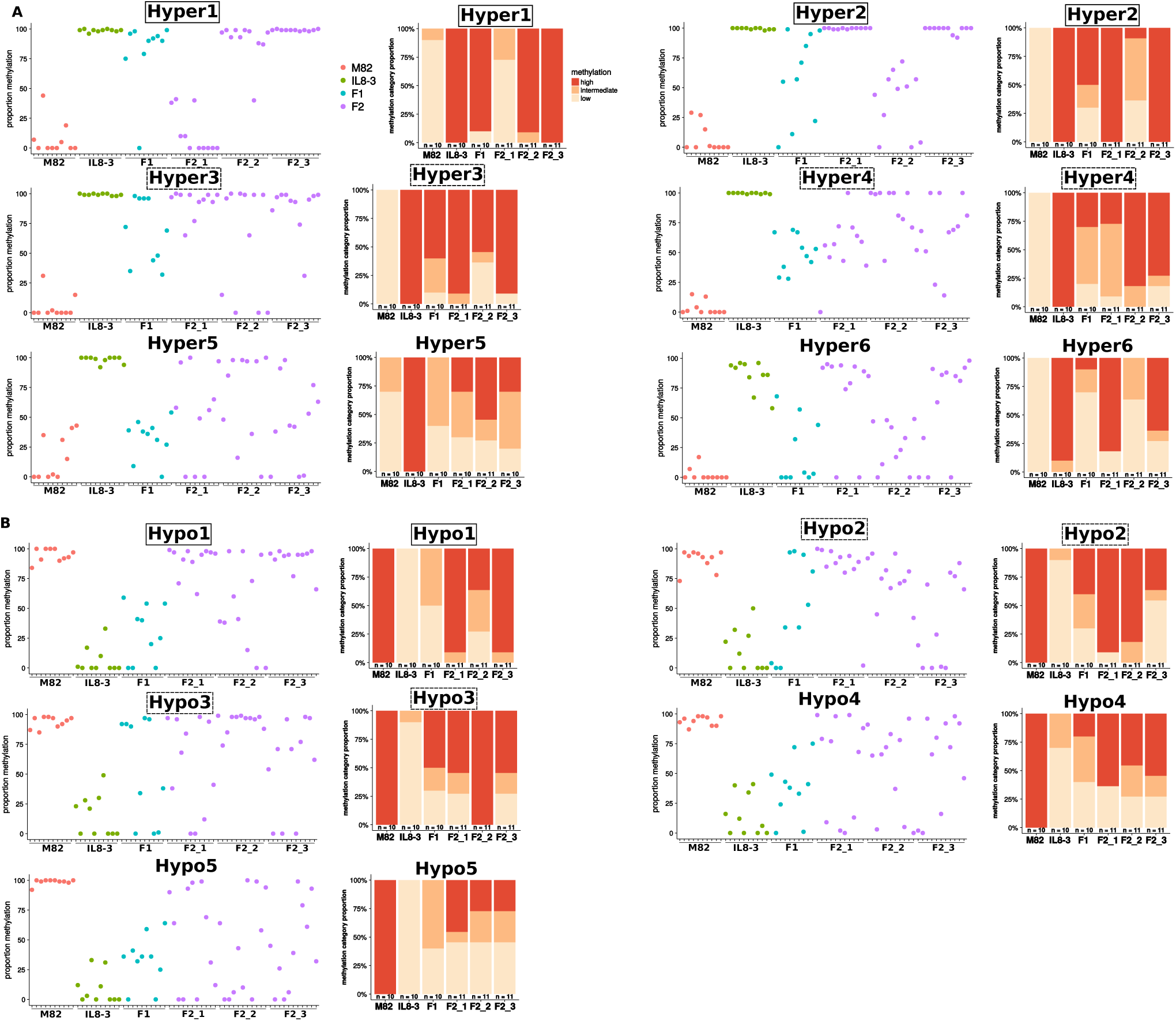
Validation of paramutation by McrBC. (A) Hypermethylated DMRs. (B) Hypomethylated DMRs. A solid frame indicates strong evidence for paramutation, and a dashed frame partial evidence. For each region, the left hand panel shows the results of methylation analysis by McrBC for individual plants. This information is collated in the right hand panel with the splitting of the F2s according to their F1 parent. Low methylation: ≤ 33%. Intermediate: > 33% and ≤ 66%. High: > 66%.

It is clear, however, that the DNA methylation marks at these loci are not completely stable. At Hyper1 and Hyper2, for example, there were F1s with very little methylation, suggesting that there must have been loss of methylation even from the IL8-3 allele. It is likely therefore that there are two oppositely acting mechanisms influencing the methylation status of these loci: a paramutation-like event leading to *de novo* establishment of DNA methylation and a second process leading to removal of DNA methylation. The net effect of these processes is likely to account for the extent to which the pattern of DNA methylation deviates from Mendelian ratios in these F1 and F2 plants. However once established in the F1, methylation may be fairly stable: F2 progeny from two out of three F1s were consistently highly methylated for Hyper1 and Hyper2.

In addition to these four loci (Hyper1–4, Fig. 7A) we also investigated inheritance at five other loci that were hypermethylated in the IL1-1, IL2-5 and IL8-3 (Fig. S3 and Table 1). At Hyper5 and Hyper6 there was no conclusive evidence for paramutation-like behaviour (Fig. 7A) and we conclude either that these are conventional Mendelian loci or that the instability of the methylation mark in the progeny of M82 × IL8-3 offset the effects of the paramutation-like mechanism. At Hyper7 and Hyper8, high methylation in some M82 plants as assessed by McrBC does not allow us to conclude that the lack of F2s with low methylation reflects non-Mendelian segregation (Fig. S3). Methylation in IL8-3 at Hyper9 was variable, and there was no evidence for paramutation-like interactions (Fig. S3).

#### Hypomethylated DMRs

We also tested inheritance of 8 hypomethylated DMRs in IL1-1, IL2-5 and IL8-3 (Hypo1–8, Fig. 7B and Fig. S3). For one locus (Hypo1), there was a high proportion of the hypermethylated allele in the F2 (from two F1s in particular) although in the F1 the trend was towards medium-to-low methylation. These proportions indicate that methylation of this locus is subject to instability but that the hypermethylated M82 epiallele has some paramutation-like activity.

Similarly at Hypo2 andHypo3, there was evidence for contrasting dynamics of methylation: 4 and 5 (out of 10) F1s were highly methylated, while 3 in each case had very low methylation (Fig. 7B). Overall the proportions of lowly methylated F2s did not significantly depart from Mendelian ratios, but the F1 of origin had clear effects, with progenies of some F1s homogeneously hypermethylated. These results argue for paramutation-like activity as well as instability of the hypermethylated M82 epialleles.

At Hypo4 and Hypo5 there was no evidence for non Mendelian segregation (Fig. 7B), while for the remaining three loci (Hyper6–8, Fig. S3), McrBC estimates of methylation in the parents were too variable to interpret the methylation levels of their progeny.

In summary, of the 17 DMRs tested in detail, 7 showed at least partial evidence of a heritable gain of methylation upon crossing (Table 1). Because the methylation differences were observed in three independent introgression lines, it is unlikely that they are due to a *trans* effect from the introgressed *S. pennellii* region and a more likely explanation is that, when methylation is gained in the M82 × IL8-3 F1s and F2s there is an epigenetic cause.

## Discussion

### Origins of DNA methylation differences: paramutation-like mechanisms versus spontaneous epigenetic variation

The main aim of this study was to explore the epigenetic changes associated with wide cross hybridisation of M82 × *S. pennellii* through the characterisation of *H06* and other DMRs in at least three different ILs relative to the parental lines. In principle these shared DMRs, as with DMRs in hybrid progeny of maize (18), rice (19) and *Arabidopsis thaliana* (12, 20), could be due to spontaneous epimutation (21, 22) or genetic differences in seedstocks (e.g. a transposon insertion) (23, 24) as well as to genetic or epigenetic interactions between the parental genomes. For many of the DMRs shared by the three ILs under study, we cannot say which of these explanations is likely to apply. For *H06*, however, we could reproduce its epimutation whenever we made the M82 × *S. pennellii* cross and paramutation in backcrosses of IL8-3 with M82 occurred at at high frequency, suggesting that the epigenetic state of the ILs at *H06* is due to the hybridisation process.

To explain hybrid-induced paramutation it could be that there are genetic or epigenetic differences between the parental genomes. A genetic mechanism might apply if the two genomes differ by insertion or deletion mutation at the affected locus. In such a scenario it could be, by analogy with epigenetic marks in *Neurospora crassa* that the transition is triggered by unpaired DNA during meiosis of the F1 (25). The difference between the plant and fungal systems in the properties of the epigenetic mark: in *N. crassa* it would be a standard heritable mark with Mendelian inheritance whereas in tomato it would be a heritable mark that can transmit between alleles.

This requirement for unpaired DNA would account for the delayed onset of the paramutation-like process of *H06* until the meiosis in the reproductive phase of the F1 (Fig. 3). An alternative epigenetic explanation could invoke sRNAs from one of the interacting alleles that, in the parental line, are at too low an abundance to trigger paramutation. If the second allele has characteristics that favour production of secondary sRNA, for example repeats (5), then the formation of the hybrid would trigger a high level of sRNA and RdDM. The well established feedback loops of the RdDM process (26) would ensure that, once established at one allele, the amplified RdDM is stable and has the epigenetic drive characteristic of paramutation in subsequent generations.

These hypotheses are not mutually exclusive because both RdDM and meiotic silencing of unpaired DNA are dependent on an RdRP (25, 27). The involvement of RdDM is supported by several lines of evidence that link small RNAs and in particular 24-nt sRNAs with paramutation-like events in tomato in addition to the findings from other species (5, 11–13). At *H06*, for example, the timing of the paramutation-like process correlates with the timing or sRNA production (Fig. 3) and in VIGS the viral sRNAs produced could mediate (although inefficiently) the stable methylation of *H06* and the onset of paramutation-like properties (Fig. 4). In addition to *H06*, out of four loci where the hypermethylated epiallele was associated with upregulated sRNAs, three showed evidence of paramutation-like behaviour (Hyper1, Hyper2 and Hypo3, Fig. 7 and Table 1).

There are, however, epialleles with paramutation-like characteristics (Hyper3, Hyper4, Hypo1 and Hypo2, Fig. 7) that are not associated with differential small RNAs. It remains possible that RNA-independent mechanisms are involved although some of these examples may be due to sRNA production and gain of methylation in tissues or developmental stages that are not well represented in the samples used for this analysis. Conversely, not all epialleles with differential sRNA have distinct epigenetic marks or, even if they do, are paramutagenic. Clearly paramutation requires additional, as yet unknown, factors beyond those associated with heritable epigenetic marks.

### Stability of epialleles and penetrance of paramutation-like effects

For one locus with two epialleles (unmethylated and methylated), there are four possible states in a non-mosaic diploid organism: ‘ unmethylated/unmethylated’, ‘ unmethylated/methylated’, ‘ methylated/unmethylated’, ‘ methylated/methylated’ (Fig. S4). Local positive feedback loops in the maintenance of methylation (26, 27) or non-methylation (28) contribute to the stability of each state, while spontaneous epimutations and epiallele interactions in epi-heterozygotes trigger state transitions and distortions in the expected Mendelian ratios (Fig. S4).

Epialleles associated with paramutation in maize can be highly stable as with the *b1* locus, or unstable and revert to the active state, as with the *pl1* locus (29). Similarly, with the paramutation-like loci in tomato there are varying degrees of stability. For some loci there were F1 plants without methylation (*H06*, Hyper1, Hyper2, Hypo1, Hypo2, Hypo3, Fig. 7), indicating a degree of instability of the methylated epiallele. Currently there are no clear mechanisms to explain the loss of methylation in an epi-heterozygote, apart from a dilution by a factor two of the sRNAs that could weaken a RdDM positive feedback loop acting in trans.

The penetrance of the paramutation-like interaction (transition to the ‘ methylated/methylated’ state) also varied: while 9/10 F1s had high methylation at Hyper1, only 5/10 were highly methylated at Hyper2 (Fig. 7). It is therefore crucial to garner information about stability and paramutagenicity of a large number of epialleles in order to identify their determinants. However we note that a binary depiction of the epigenetic state (e.g. methylated/unmethylated) is not complete: while *H06*^*IL ′*^ is very stably methylated, the paramutated *H06*^*IL′*^ can revert when backcrossed to M82 (Fig. 2).

### Frequency of paramutation in tomato hybrids

Paramutation with perfect efficiency will be fixed very rapidly in a population and will therefore be difficult to detect. Using a wide cross such as *Solanum lycopersicum* cv. M82 × *Solanum pennellii* LA0716 increases the likelihood of detecting highly penetrant paramutation events. We based our genome-wide search for paramutation based on the characteristics of *H06* paramutation, which is associated with a gain of methylation in all sequence contexts and abundant sRNAs in multiple ILs at the seedling stage. We only found one other locus, Hyper1, that shared all of these characteristics, and this locus also displayed paramutation-like properties. However segregation analyses (Fig. 7) revealed that loci without differential sRNAs or without hypermethylation in all contexts could also engage in paramutation-like behaviours. Unexpectedly, loci that were hypomethylated in the ILs could also gain methylation upon crossing to M82.

These findings suggest that paramutation-like interactions are in fact common; occurring at hundreds of loci in a M82 × IL8-3 cross. However fully penetrant paramutation is rare: primary paramutation at *H06* was the most penetrant, but it displayed weak secondary paramutation (Fig. 2); and the penetrance of epigenetic changes and their inheritance in the seven validated loci (from seventeen tested) was incomplete. Therefore we conclude that full penetrance is rare and the IL crossing scheme may be too stringent (three consecutive backcrosses) to identify many traces of paramutation from the initial M82 × *S. pennellii* cross. Further studies aiming at assessing the prevalence of paramutation will need to be powered to detect incomplete penetrance.

### Effects on gene expression

DNA methylation is primarily associated with the silencing of repeated sequences. Consistently earlier examples of paramutation are often linked with transposable elements (5). Thus, as a mechanism of identity-based silencing, paramutation is thought to contribute to silencing multiple copies of transposable elements throughout the genome, and differential methylation may only occasionally also affect gene expression. We found no evidence for differential expression of the genes surrounding *H06* in the seedling expression data set. However *H06* is a target of the RIN (RIPENING INHIBITOR) transcription factor (30) and its surrounding genes are most highly expressed in the ripening fruit (31) (Fig. S5), so its differential methylation may have an effect in this particular tissue. For three of the validated DMRs that engaged in paramutation-like interactions (Hyper3, Hyper4 and Hypo3), the neighbouring genes were differentially expressed in the introgression lines compared to M82 (Fig. S2).

Generating novel epialleles can be a source of phenotypic diversity (32, 33). If the epialleles are stable enough and their epigenetic state can be efficiently controlled, epigenetic drives via paramutation may be used to improve crops.

## Conclusions

In this study we show that interactions between natural tomato epialleles are relatively common, can lead to heritable differences in methylation, and occur mostly in one direction (gain of methylation) while stochastic loss of methylation may act as a counterbalance. There were however variations between pairs of epialleles in timing, penetrance and heritability of the gain of methylation. We propose that interactions between epigenomes should be viewed as a continuum, of which fully penetrant and fully heritable paramutation is an extreme manifestation. Taking into account the partial penetrance and stochasticity of epigenetic interactions will be crucial to properly investigate transgenerational inheritance.

## Methods

### Plant material

Tomato (*Solanum lycopersicum*) cv M82, *S. lycopersicum* Micro-Tom, *Solanum pimpinellifolium* and *Solanum pennellii* plants were raised from seeds in compost (Levington M3) and maintained in a growth room at 23°C with 16/8 h light/dark periods with 60% relative humidity, at a light intensity of 150 *μ*mol photons m^−2^.s^−1^. Unless otherwise indicated, *S. pennellii* refers to accession LA0716 that was used to generate the introgression lines (14).

### DNA extraction

DNA from leaf (2-week-old seedlings), mature flowers or pollen was extracted with the Puregene kit (QIAgen) following manufacturer’s instructions.

### Genotyping

IL8-3 was genotyped based on the TG510 marker: the sequence was amplified with primers TG510 fw and TG510 reverse, and digested with AluI. The M82 sequence is cleaved in a 133-bp and a 245-bp fragment, while the introgressed sequence from S. pennellii remains intact (378 bp). The *H06* sequence from M82 was differentiated from *S. pennellii* and *S. pimpinellifolium* by digesting *H06* amplicons with HpaI. The *S. pennellii* and *S. pimpinellifolium* are digested into two fragments, 260 and 210 bp in length, while the M82 sequence remains uncut. Even brightness of the *H06* genotyping bands for F1s on an agarose gel indicate that McrBC-qPCR results should not be distorted by differences in DNA amplification efficiency between alleles.

### RNA extraction

Total RNA samples were prepared from 100 mg of tissue using TRIzol (LifeTechnologies).

### VIGS

*H06* genomic insert was cloned into the binary TRV RNA2 vector using the KpnI and XhoI restriction sites of the multiple cloning site as described previously (16, 34). Cotyledons of tomato seedlings were agro-infiltrated 10 d after sowing with a 1:1 mixture of *Agrobacterium tumefaciens* (strain GV3101:pMP90+pSOUP) carrying TRV RNA1 and RNA2 at OD_600_ = 1.5.

### Viral load quantification

Quantification of viral load was performed with Precision One-Step qRT-PCR (Primerdesign) on 15 ng RNA per well and normalised with *TIP41*.

### sRNA-Seq

sRNAs from leaf (2-week-old seedlings), flower and pollen of M82, *S. pennellii* LA0716 and their F1 were cloned from 10 *μ*g total RNA using the Illumina TruSeq Small RNA cloning kit and libraries were indexed during the PCR step (12 cycles) according to the manufacturer’s protocol. Gel size-selected, pooled libraries were sequenced on a HiSeq 2000 50SE. Padubiri Shivaprasad prepared Micro-Tom and *S. pimpinellifolium* sRNA libraries from the aerial part of two-week-old seedlings according to (15). Sequences were trimmed and filtered with Trim Galore! (with the adapter parameter ‘ -a TGGAATTCTCGGGTGCCAAGG’). Mapping to *H06* (*S. pennellii* and M82 sequences) was performed with Bowtie 1.1.1 (35) without mismatches (options ‘ -v 0 -a’).

sRNA counts on *H06* in Fig. 1 were obtained by mapping sRNA libraries from (15) to Heinz genome SL2.50 with Bowtie 1.1.1 without mismatches (options ‘ -v 0 -a’), and extracting the counts for the interval “SL2.50ch08:54487325-54487842". We found that there was a mistake in the labelling of the libraries deposited in (15): the libraries labelled IL1-1, IL2-5,IL8-1-1, IL8-1-D, IL8-1-3, IL8-2, IL8-2-1, IL8-3 and IL8-3-1 actually correspond to IL8-3-1, IL8-3, IL8-2-1, ILH8-2, IL8-1-5, IL8-1-D, IL8-1-1, IL2-5 and IL1-1. For convenience the correctly labelled sRNA libraries for the introgression lines used in this study (IL1-1, IL2-5 and IL8-3) are included in the GEO data set.

To find differential sRNA loci between M82 and ILs, sRNA libraries from IL1-1, IL2-5, IL8-3 and two M82 seedlings (from (15)) were mapped and clustered on Heinz genome SL2.50 using ShortStack v3.3.3 (36) with default parameters. sRNA counts on the defined loci were analysed with DESeq2 v1.8.1 (37).

### Gene expression analysis

Transcript counts for two-week-old IL1-1, IL2-5, IL8-3 and M82 seedlings from (17) were analysed with DESeq2 (37) with the design factors ‘ condition’ (sun or shade), ‘ experimental batch’, and ‘ genotype’ (either IL or M82). Genes were considered differentially expressed between genotypes when the adjusted p-value was for the ‘ genotype’ factor was *<* 0.05.

### Analysis of DNA methylation

#### McrBC

Analysis of methylation by McrBC was performed as previously described in (16).

#### Sanger Bisulfite

For Sanger bisulfite sequencing, 450 ng of DNA was bisulfite-converted with EZ DNA Methylation-Gold Kit (Zymo Research) and amplified with primers specific to the region and the Kapa Uracil+ HotStart DNA polymerase (Kapa Bioscience). Amplification products were size selected on a 1.5% agarose gel and gel extracted with the QIAquick gel extraction kit (QIAgen), A-tailed following the same protocol as for the library preparation (below), and cloned into pGEM-T easy (Promega). Sequences aligned with MUSCLE were then analysed with CyMATE (38) and PCR-duplicates were removed.

#### MethylC-Seq

Bisulfite library preparation was performed with a custom protocol similar to ref (39). 1.2 *μ*g DNA was sonicated on a Covaris E220 to a target size of 400 bp and purified on XP beads (Ampure, ratio 1.8). DNA was end-repaired and A-tailed using T4 DNA polymerase and Klenow Fragment (NEB) and purified again using XP beads (ratio 1.8x). Methylated Illumina Y-shaped adapters for paired-end sequencing were ligated using Quick-Stick Ligase (Bioline). 450 ng of purified (ratio 1.8x), adapter-ligated DNA was bisulfite-converted using EZ DNA Methylation-Gold Kit (Zymo Research) according to manufacturer’s instructions. DNA was barcoded using 12 cycles of PCR amplification with KAPA HiFi HotStart Uracil+ Ready Mix (Kapabiosystems) with PE1.0 and custom index primers (courtesy of the Sanger Institute). Duplicate libraries for M82, IL1-1, IL2-5 and IL8-3 (leaves of two-weekold seedlings) were sequenced in paired-end mode. Reads were trimmed and filtered with Trim Galore! v0.4.2 (default parameters), then mapped on Heinz genome SL2.50 using Bismark v0.17.0 (40) (first in paired-end mode with options ‘ –score-min L,0,-0.2 -p 4 –reorder –ignore_quals –no-mixed –no-discordant –unmapped’, then unmapped read1 was mapped in single-end mode with the same quality parameter ‘ -N 1’). Reads were deduplicated with ‘ bismark_deduplicate’ and methylation calls were extracted using Bismark ‘ methylation_extractor’ (with options ‘ -r2 2’ for paired-end reads). Based on Bismark’s cytosine report, methylated and unmethylated counts for cytosines of both strands were pooled into 300-bp bins sliding by 200 bp and separated by context (CG, CHG, and CHH). Bins with fewer than 10 counts or with coverage exceeding the 99^*th*^ percentile in one or more libraries were excluded from the analysis. We also excluded the introgressed regions: ch01 1 – 86 Mb, ch02 44 – 54 Mb and ch08 61 Mb – 65866657 bp (end of chromosome). Each introgression line was then compared to M82 by fitting a logistic regression on the methylated and unmethylated counts (R glm with ‘ family = binomial’). The p-values were subjected to a Benjamini-Hochberg correction to control the false discovery rate at 5%. Differentially methylated bins were further filtered by imposing thresholds on the absolute difference in methylation (average of replicates) between M82 and the introgression line: a difference of 25% in the CG context, 20% in CHG and 10% in CHH. DMRs were constructed by merging overlapping differentially methylated bins. We included the region Hyper2 into the McrBC validation analysis despite it being inside the introgressed region in IL2-5, because it was hypermethylated in CG, CHGand CHH contexts and produced abundant sRNAs in IL1-1 and IL8-3, thus making it a likely candidate for paramutation. Chloroplast DNA was used as a control for bisulfite conversion efficiency, and sequencing statistics are collected in additional file 3.

#### Oligonucleotides

Please refer to additional file 1.

## DATA AVAILABILITY

The datasets generated and analysed during the current study are available in the GEO repository under accession GSE97247.

## AUTHOR CONTRIBUTIONS

QG and DCB designed the research and wrote the manuscript, QG performed the research.

## FUNDING

This work was supported by the European Research Council Advanced Investigator Grant ERC-2013-AdG 340642, the Balzan Foundation, the Biotechnology and Biological Sciences Research Council and the Frank Smart Studentship. D.C.B. is the Royal Society Edward Penley Abraham Research Professor. The funding bodies had no roles in the design of the study, in the collection, analysis, interpretation of data, or in the writing of the manuscript.

### ACKNOWLEDGEMENTS

The authors would like to thank Shuoya Tang and Mel Steer for horticultural assistance, Aleksandar Ivanov for his work on sRNA libraries, Padubiri Shivaprasad for the Micro-Tom and *S. pimpinellifolium* sRNA libraries, Bruno Santos, Felix Krueger, and Sebastian Müller for bioinformatics advice.

## Additional Files

Additional file 1 — Oligonucleotides used in this study. Tsv format.

Additional file 2 — H06 sequences for *S. lycopersicum* cv. M82, *S. pimpinellifolium* and *S. pennellii* LA0716. Fasta format.

Additional file 3 — Summary of MethylC-seq data. Tsv format.

Additional file 4 — DMRs shared between introgression lines. Tsv format.

Additional file 5 — DMRs shared between introgression lines and hypermethylated in all cytosine contexts. Average proportion of methylated cytosines between two replicates, for each context and genotype. Tsv format.

Additional file 6 — DMRs shared between introgression lines and hypomethylated in all cytosine contexts. Average proportion of methylated cytosines between two replicates, for each context and genotype. Tsv format.

## Supplementary Figures

Supplementary figure 1 — Genomic position, small RNAs and DNA methylation of the IL DMRs Hyper1–9 and Hypo1–8. The selected DMR is highlighted in red, the plotted region includes 2 kb upstream and downstream. Genes: ITAG2.4 gene models. Repeats: RepeatMasker annotation. Hypomethylated regions shared between the three introgression lines are annotated in blue, and hypermethylated regions in red. sRNAs: coverage of sRNAs in seedlings. Methylation: percentage of methylated cytosines in 100-bp regions in each context (two replicates per genotype).

Supplementary figure 2 — Differential expression of genes whose promoters overlap shared IL DMRs selected for segregation analysis. Hypermethylation of Solyc09g064640’s promoter (Hyper3 and Hyper5) is associated with decreased expression in introgression lines. Solyc04g025030 is associated with Hyper4, and Solyc06g075840 with Hypo3. Normalised counts in seedlings (40 M82 and 30 ILs, data from (17)).

Supplementary figure 3 — Segregation of DNA methylation patterns by McrBC. These regions show variable methylation in the parental controls M82 and IL8-3. (A) Hypermethylated DMRs Hyper7–9. (B) Hypomethylated DMRs Hypo6–8. For each region, the left hand panel shows the results of methylation analysis by McrBC for individual plants. This information is collated in the right hand panel with the splitting of the F2s according to their F1 parent. Low methylation: ≤33%. Intermediate: > 33% and ≤ 66%. High: > 66%.

Supplementary figure 4 — Schematic of possible epiallelic transitions for one locus in a diploid state with unmethylated (white) and methylated (black) epialleles. There are four possible epigenetic configurations. Recurrent (curved) arrows represent the propensity to conserve the current state, from the action of local positive feedback loops maintaining methylation/unmethylation. Transition (straight) arrows between states represent spontaneous epimutations and, when starting from epigenetically heterozygous states, paramutation-like interactions. The epigenetic state of a genome may be seen as the result of such a stochastic process at each cell/organism generation.

Supplementary figure 5 — Potential role of H06 in fruit ripening. Expression profiles across tissues of the two genes immediately downstream of *H06*, Solyc08g066020 (A) and Solyc08g066030 (B). Their expression peaks in mature green fruit. Data from the eFP Browser (31). (C) Demethylation of the 3’ end of *H06* during fruit ripening correlates with the binding of the RIN transcription factor at this locus. RIN binding, demethylation and ripening are compromised in the *rin* (ripening inhibitor) and *cnr* (colorless non-ripening) mutants. Data from (30).

